# Mutualistic interactions with phoretic mites *Poecilochirus carabi* expand the realised thermal niche of the burying beetle *Nicrophorus vespilloides*

**DOI:** 10.1101/590125

**Authors:** Syuan-Jyun Sun, Rebecca M. Kilner

## Abstract

Mutualisms are so ubiquitous, and play such a key role in major biological processes, that it is important to understand how they will function in a changing world. Here we test whether mutualisms can help populations to persist in challenging new environments, by focusing on the protective mutualism between burying beetles *Nicrophorus vespilloides* and their phoretic mites (*Poecilochirus carabi*). Our experiments identify the burying beetle’s fundamental thermal niche and show that it is restricted by competition with blowfly larvae at higher and lower temperatures (within the natural range). We further demonstrate that mites expand the burying beetle’s realised thermal niche, by reducing competition with blowflies at lower and higher temperatures, thereby enabling beetles to produce more offspring across a wider thermal range. We conclude that mutualisms can play an important role in promoting survival under novel and adverse conditions, particularly when these conditions enhance the performance of a common enemy.

## Introduction

Mutualisms are defined as mutually beneficial interactions between species (Bronstein 2015). They are ubiquitous to all life on earth and key to structuring biological processes from community dynamics to ecosystem function (Bronstein 2015). Nevertheless, mutualistic interactions can be unstable on an ecological timescale and are prone to sliding into more antagonistic relationships (Sachs & Simms 2006; Anderson & Midgley 2007; Hoek *et al.* 2016), depending on the wider ecological or physical environments in which they are embedded (Thrall *et al.* 2007; Jovani *et al.* 2017). Consequently, there is considerable interest in determining how mutualisms are likely to respond to environmental degradation (Kiers *et al.* 2010), and to rising temperatures in particular (Doremus *et al.* 2018).

Previous work has emphasized the vulnerability of mutualisms to environmental change. Increased temperatures cause some mutualistic interactions to break down because partner species become phenologically mismatched (Rafferty *et al.* 2015; Renner & Zohner 2018) or because they sustain different levels of thermal tolerance (Barton & Ives 2014; Fitzpatrick *et al.* 2014; Sagata & Gibb 2016; Doremus & Oliver 2017; Zhou *et al.* 2017). Nevertheless, other partnerships between species can withstand exposure to higher temperatures (e.g. Zhou *et al.* 2017), and then it becomes harder to predict whether the mutualism will persist or degrade.

The effects of rising temperatures are especially difficult to predict for protective mutualisms. In this class of mutualism, one species provides a resource for its partner in exchange for defence from parasites or predators (Ostlund-Nilsson *et al.* 2005). The effect of rising temperatures on the mutualists’ natural enemies is key to predicting the mutualism’s fate. For example, if the enemy species succumbs to heat stress before the protective partner species, and the mutualism is rendered redundant, then the protective mutualist might then become an antagonist instead (e.g. (Okabe & Makino 2008)). Conversely, if the enemy species and mutualists all thrive as temperatures rise then the protective mutualism might be strengthened, through a greater need for the defensive service provided by the protective partner (Cheney & Côté 2005). The fate of the mutualism can then be described in terms of niche theory (Johnson 2015). If an enemy species thrives at higher (or lower) temperatures then a species’ realised niche might become even smaller than its fundamental niche. However, a protective mutualist can potentially counteract these effects and enable an organism to expand its realised niche, even when the enemy species performs better at higher (or lower) temperatures. Whether this ever happens, however, is largely unknown.

Here we determine the effect of a putative protective mutualism between burying beetles *Nicrophorus vespilloides* and their phoretic mites *Poecilochirus carabi*, on the burying beetle’s realised thermal niche. Both species breed on the body of a small dead vertebrate like a mouse or a songbird. The burying beetle serves the mite by transporting it to this essential breeding resource, and by dispersing the next generation of mites at the end of each reproductive bout. Whilst they are onboard the beetle, the mites are benign passengers. The putative protective mutualism arises during reproduction on the corpse. At this point, the mites potentially serve the burying beetle by consuming eggs and larvae of rival blowflies (Calliphoridae): this is how they behave when carried by congeneric burying beetles *N. orbicollis* and *N. tomentosus* (Springett 1968, Sloan Wilson 1983). It is unclear whether this form of protective mutualism exists between *N. vespilloides* and *P. carabi.* However, these two species are known to be antagonists when blowflies are not present. At high densities, mites sometimes attack beetle larvae directly (Wilson & Knollenberg 1987; De Gasperin & Kilner 2015) and compete with them for carrion (Wilson & Knollenberg 1987; De Gasperin *et al.* 2015). Therefore if blowflies disappear with rising temperatures, the interactions between mites and burying beetles will become more antagonistic.

We used a combination of field and laboratory experiments to address three related questions: 1) Do *P.carabi* protect breeding burying beetles *N. vespilloides* from competition with blowfly larvae? 2) Does the effectiveness of this putative protective mutualism vary with temperature? 3) Specifically, does *P. carabi* increase the realised thermal niche of the burying beetle?

## Material and methods

### Study system

Burying beetles (*N. vespilloides*) use small vertebrate carcasses as their sole breeding resource. Both males and females convert the carcass into an edible carrion nest, by removing any fur or feathers, rolling it into a ball, smearing it with antimicrobial exudates (Cotter & Kilner 2010), and burying it underground. During carcass preparation, eggs are laid in the surrounding soil and hatch within 3-4 days. The larvae feed themselves on the edible nest and also fed by both parents. Approximately 4-5 days after hatching, the larvae disperse away from the scant remains of the carcass to pupate. Adult burying beetles carry up to 14 species of phoretic mites (Wilson & Knollenberg 1987). The *P. carabi* species complex is the most salient and common of these mite species. In natural populations, 84.4% (1156 out of 1369) of trapped adult *N. vespilloides* carry 0-20 *P. carabi* mites (Sun *et al.* 2019).

A dead body is a rare “bonanza resource” (Scott 1998) generating competition within and among species for the resources upon it. Blowflies (Calliphoridae) are a particular competitive threat for burying beetles. They can more rapidly locate the newly dead and start to lay eggs within minutes of arriving on the dead body (Bornemissza 1957; Payne 1965; Matuszewski *et al.* 2010). Their larvae also develop rapidly, quickly consuming resources on the corpse. Nevertheless, their breeding success is modulated by temperature. Previous work shows that at lower temperatures blowflies are less abundant on carrion, and that they develop more slowly and they have lower reproductive success (Wall *et al.* 1992; Sun *et al.* 2014).

### Burying beetles and phoretic mites in Madingley Wood

Fieldwork was carried out at Madingley Woods in Cambridgeshire UK, an ancient woodland of mixed deciduous trees near the Sub-Department of Animal Behaviour, University of Cambridge, (Latitude: 52.22730°; Longitude: 0.04442°). We assessed the density of *N. vespilloides* and the mite *P. carabi* by setting Japanese beetle traps, baited with ∼ 30 g fresh mice, from June to October, 2016-2017. Ambient air temperature was recorded locally at 1 h interval using an iButton temperature data logger (*n* = 8; DS1922L-F5#, Maxim Integrated Products, Inc.), which was suspended alongside each trap at 1 m above the ground. Traps were checked daily to determine when the beetles first located the dead body. We found the mean ± S.E.M time to discovery was 3.42 ± 0.77 days. Each trap was emptied every two weeks, and rebaited fresh mouse carcass. At this point, we counted the total number of *N. vespilloides* and the number of *P. carabi* carried by each individual beetle. Beetles were temporarily anaesthetized in the lab using CO_2_ and mites were then detached with a fine brush and a tweezer. Field-caught beetles, mites, and blowfly larvae collected from the traps were used to establish laboratory colonies (see Supplementary Information for full details of their husbandry).

### Field experiments

To assess whether mites are protective mutualists of the burying beetle, we investigated how they affected burying beetle reproductive success in the field, in Madingley Woods. These experimental breeding events involved opportunistic cobreeding by the blowflies that were present naturally in the woods. We recorded ambient temperature during each experiment by using iButton temperature data loggers placed at 1 m above ground at 1 h intervals throughout. Breeding events were established at 20 different sites (see Fig. S1), each separated by approx. 30 m from the nearest neighbouring site. Each site was used more than once. The set up for each breeding event is shown in Fig. S2. A 8-16 g (12.40 ± 0.15 g) mouse carcass was placed on the compost and left for three days, to simulate the average time taken by beetles to locate a carcass in the field (see above). We then added a pair of burying beetles from the laboratory colony. We also added mites at one of three different densities: 0, 10, or 20 mite deutonymphs (*n* = 66, 68, and 61 respectively for these three treatments). At Madingley Woods, 113 out of 172 (65.7%) wild-caught *N. vespilloides* carried 0 – 92 mites per beetle (median = 5). Therefore these manipulations of mite density fall within the natural range.

Each experiment was terminated when the beetle larvae dispersed or when the dead body was completely consumed by blowfly larvae. At this point we measured components of beetle fitness (number of beetle larvae), blowfly fitness (number of blowfly larvae), and mite fitness (number of dispersing mite deutonymphs on adult beetles).

### Laboratory experiment 1

We repeated the experiment in a lab setting so that we could carry out manipulations to address two specific questions:

#### 1) Are mites in a protective mutualism with burying beetles?

Here we tested for evidence that a) blowflies depress burying beetle fitness and b) mites can counteract any such negative effects. Accordingly, we manipulated the presence/absence of blowflies and mite density, using a fully-factorial 2 (blowfly treatments) x 3 (mite treatments) experimental design. To simulate the presence of blowfly competition, we placed 30 mg (30.22 ± 0.07 mg) of newly-laid blowfly eggs onto a 7-16 g (11.13 ± 0.15 g) mouse carcass, mimicking the rapid oviposition on a freshly dead carcass by blowflies in nature (Wilson 1983). As a control, dead mice of similar size were kept free of blowflies (10.64 ± 0.15 g). In both blowfly treatments, the dead mouse was placed on the soil in a breeding box in a temperature-regulated breeding chamber for 3 days before adding the beetles, simulating the later arrival of the beetle at the carcass that is seen in nature (see above). During this time, the fly eggs were able to hatch and the blowfly larvae started to consume the carcass. The mite density treatments matched those used in the field experiment: 0, 10, or 20 mites. Mite deutonymphs were introduced to the dead mouse at the same time as the burying beetles. When the breeding bout was complete, as indicated by either the beetle larvae starting to disperse away or carcass consumption by blowfly larvae, whichever came sooner, we measured the fitness components of beetles, mites, and blowflies using the methods described above in the field experiments.

#### 2) Is the protective mutualism modulated by temperature?

The six treatments described above were staged in temperature-regulated breeding chambers (Panasonic MLR-352-PE). Each temperature treatment mimicked the 8°C diurnal temperature fluctuation that is typical for Madingley Woods, during the burying beetle’s breeding season (Fig. S3). The mean temperature for each manipulation was 11, 15, and 19°C, which matches the mean seasonal low, intermediate, and high temperatures, respectively, in Madingley Woods (Fig. S3). Each of the six treatments was carried at these three temperatures, generating 18 treatments in all. This experiment allowed us to determine the burying beetle’s fundamental thermal niche, by quantifying beetle reproductive success across a thermal gradient in the absence of blowflies and mites. It also enabled us to determine the extent to which blowflies reduce this fundamental niche, by comparing the beetle’s reproductive success across this thermal gradient, with and without blowflies. By then introducing mites, and again measuring burying beetle reproductive success across the same thermal range, we were able to determine whether or not mites expanded the beetle’s realised thermal niche. For logistical reasons, replicates of all 18 treatments were spread over four blocks, carried out in succession.

### Laboratory experiment 2

#### Effect of temperature on blowfly larval development

In a second experiment, we examined larval development, the number of dispersing larva, and the rate of carcass consumption, at the three different temperatures used in the preceding experiment (11, 15, and 19°C; *n* = 13 carcasses for each temperature treatment). This enabled us to examine how blowflies respond to temperature, independent of the actions of the mites and blowflies. Once again, we placed blowfly eggs (30.22 ± 0.09 mg) on a mouse carcass (10.74 ± 0.30 g) placed on soil in a plastic breeding box, and put the box in a temperature-controlled breeding chamber (No burying beetles or mites were added this time). Every 12 h we checked the boxes and determined the stage of blowfly larval development attained, namely 1^st^, 2^nd^, 3^rd^ instars and post-feeding. In addition, we recorded when the carcass entered the bloating stage (indicated by swelling and putrefaction). When the larvae entered the post-feeding stage, we counted them, and recorded their total mass. From these data we determined the proportion of carcass consumed, calculated as total mass of larvae divided by initial carcass mass.

### Statistical analyses

Generalised linear mixed model (GLMM) analyses were carried out in the statistical programme R 3.4.3 using the package *lme4* (Bates *et al.* 2015). Post-hoc pairwise comparisons were performed using the package *lsmeans* (Lenth 2016) if an interaction was detected. Non-significant interaction terms were dropped from the analyses before reporting the results. Full details are given in the Supplementary Information.

## Results

### Burying beetles and phoretic mites in Madingley Wood

To examine the effects of temperature on the natural population dynamics of *N. vespilloides*, we determined how population density varies with a natural gradient of temperature. We found that population density peaked at the intermediate average daily temperature (∼15°C), and gradually decreased at both higher and lower temperature extremes (temperature^2^: χ^2^= 39.55, d.f. = 1, *P* < 0.001; temperature: χ^2^= 0.00, d.f. = 1, *P* = 0.999; Fig. S4a). Also, the total number of mites per beetle decreased as temperature increased (temperature: χ^2^= 17.96, d.f. = 1, *P* < 0.001; Fig. S4b).

### Field experiments

#### Effects of mites and temperature on burying beetles

We found that the number of burying beetle larvae present at the end of larval development, as dispersal, varied with temperature but that this relationship differed among the three mite treatments (mite x temperature^2^interaction, χ^2^= 10.81, d.f. = 2, *P* = 0.004; Fig. 1). We split the dataset by mite treatment to determine exactly how this relationship changed across the mite treatments. When there were no mites present, the burying beetle’s reproductive success showed a curvilinear relationship with temperature, peaking at intermediate temperatures but falling off markedly at low and high temperatures (Fig. 1a). When burying beetles bred alongside 10 mites, temperature explained no variation in beetle reproductive success (temperature^2^: χ^2^= 0.035, d.f. = 1, *P* = 0.851; temperature: χ^2^= 0.87, d.f. = 1, *P* = 0.351; Fig. 1b). However, when burying beetles bred alongside 20 mites, the relationship between temperature and reproductive success changed again, this time dipping slightly at intermediate temperatures peaking at lower and higher temperatures (temperature^2^: χ^2^= 5.35, d.f. = 1, *P* = 0.021; temperature: χ^2^= 5.51, d.f. = 1, *P* = 0.019; Fig. 1c).

**Figure 1.**
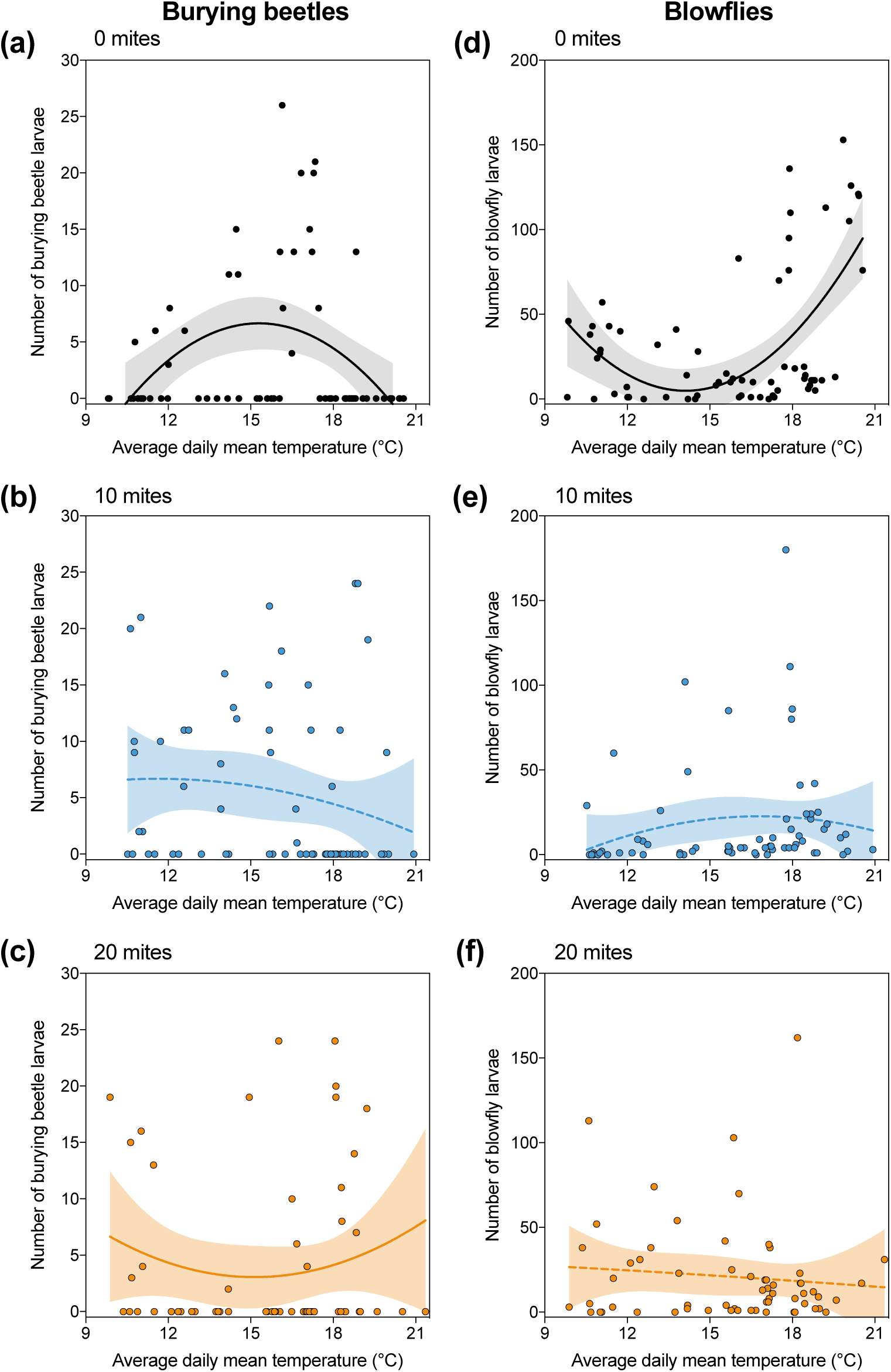
Reproductive success under field conditions in relation to ambient air temperature in (a-c) burying beetles and (d-f) blowflies, across the three different mite treatments. Shaded regions represent 95% confidence intervals, and solid and shaded lines represent statistically significant and non-significant relationships from GLMM, respectively. Each datapoint represents one breeding event.

#### Effects of mites and temperature on blowflies

In separate analyses, we investigated how blowfly reproductive success covaried with temperature and the mite treatments. Once again, we found that the relationship with temperature was different in the different mite treatments (mite x temperature^2^interaction, χ^2^= 11.53, d.f. = 2, *P* = 0.003; Fig. 1d). Again, we split the dataset by mite treatment, to see how these differences arose. We found that when no mites were present, blowflies had greatest reproductive success at higher temperatures and performed much less well at intermediate temperatures (temperature^2^: χ^2^= 9.57, d.f. = 1, *P* = 0.002; Fig. 1d), with more larvae produced at higher than lower temperature (temperature: χ^2^= 4.87, d.f. = 1, *P* = 0.027; Fig. 1d). By contrast, temperature explained much less variation in blowfly reproductive success in the 10 mite treatment (temperature^2^: χ^2^= 0.32, d.f. = 1, *P* = 0.572; temperature: χ^2^= 1.32, d.f. = 1, *P* = 0.250; Fig. 1e) and the 20 mite treatment (temperature^2^: χ^2^= 1.41, d.f. = 1, *P* = 0.236; temperature: χ^2^= 0.41, d.f. = 1, *P* = 0.521; Fig. 1f).

#### Relationship between mite reproductive success and temperature

In contrast to the beetles and blowflies, we found no evidence that mite reproductive success was affected by temperature, neither in the 10 mite treatment (temperature^2^: χ^2^= 0.48, d.f. = 1, *P* = 0.486; temperature: χ^2^= 2.44, d.f. = 1, *P* = 0.118) nor the 20 mite treatment (temperature^2^: χ^2^= 2.59, d.f. = 1, *P* = 0.107; temperature: χ^2^= 0.35, d.f. = 1, *P* = 0.552).

### Laboratory experiment 1

#### How do blowflies affect the burying beetle’s realised thermal niche?

Overall, we found that burying beetle reproductive success was affected by blowfly competition but that the magnitude of the effect depended both on mite density and temperature (fly x mite x temperature interaction, χ^2^= 76.29, d.f. = 4, *P* < 0.001; Fig. 2). To unpick these effects, we initially split the dataset by the three different mite treatments, to determine the effect of blowflies on burying beetle reproductive success at different temperatures and how that relationship was modulated by mites. The burying beetle’s fundamental thermal niche is shown in Figure 2d. In the absence of mites, blowflies reduced the beetle’s realised niche (Figure 2a). Blowflies depressed burying beetle reproductive success, but only at lower and higher temperatures (blowfly treatment x temperature treatment, χ^2^= 120.30, d.f. = 2, *P* < 0.001). Blowflies caused a significant reduction in the number of dispersing beetle larvae at low (*post-hoc* comparison, with v. without blowflies: *z* = 4.95, *P* < 0.001) and high (*post-hoc* comparison, with v. without blowflies: *z* = 11.69, *P* < 0.001) temperatures. When 10 mites were added, similar deleterious effects of blowflies were detected and they too were temperature dependent (blowfly treatment x temperature treatment, χ^2^= 41.18, d.f. = 2, *P* < 0.001). This time blowflies caused a significant drop in beetle reproductive success only at higher temperatures (*post-hoc* comparison, with v. without blowflies: *z* = 5.95, *P* < 0.001). When 20 mites were added, we again found deleterious effects of blowflies but they were temperature dependent in a different way (blowfly treatment x temperature treatment, χ^2^= 9.75, d.f. = 2, *P* = 0.008). This time, blowflies reduced beetle reproductive success only at intermediate temperatures (*post-hoc* comparison, with v. without blowflies: *z* = 3.84, *P* < 0.001).

**Figure 2.**
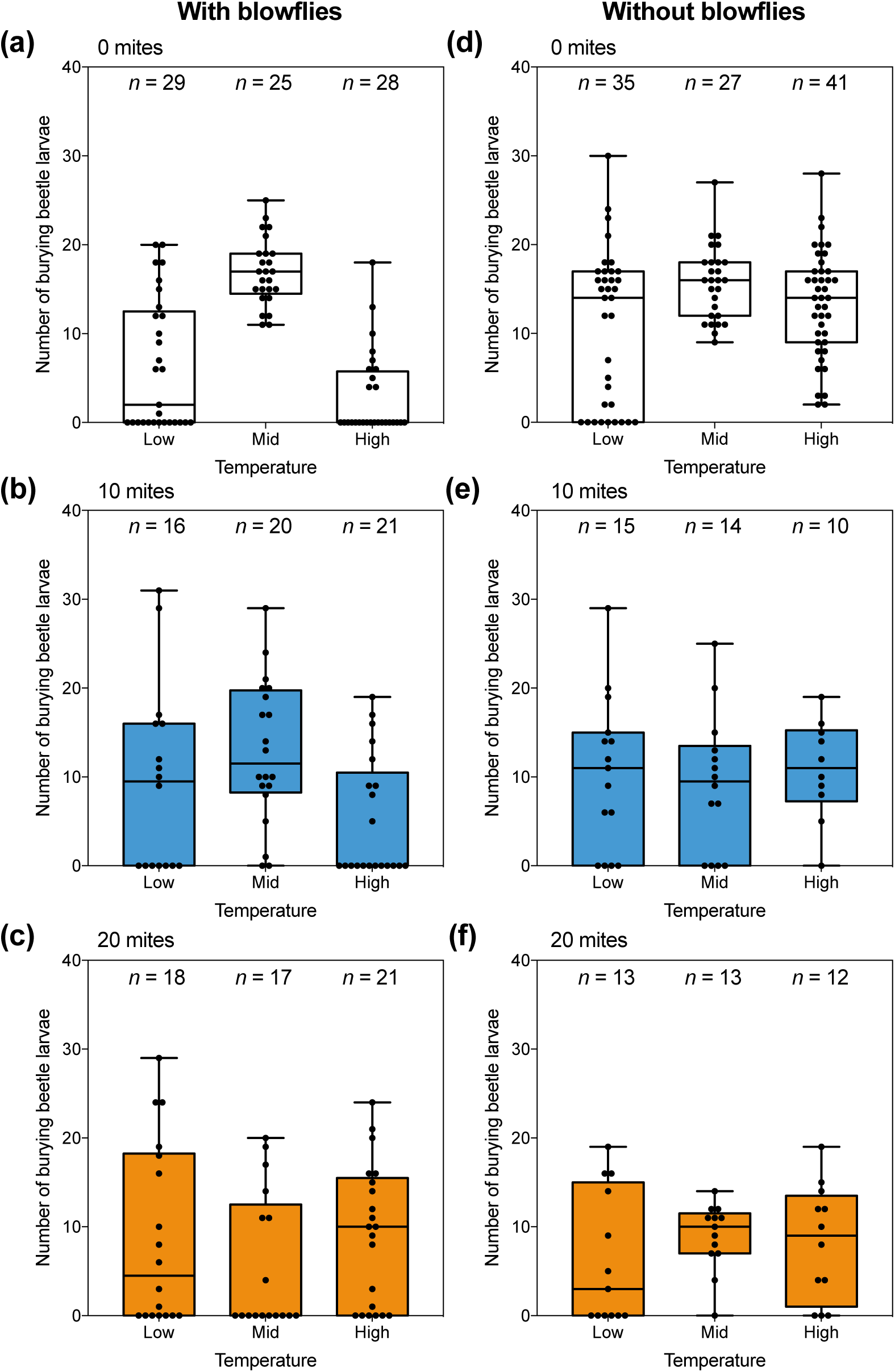
Burying beetle reproductive success under lab conditions in relation to ambient air temperature in the incubator, with and without blowflies, and across the three different mite treatments. Sample sizes are as indicated above each boxplot. The boxplots show the median, inter-quartile range, and the range of data. Each datapoint represents one breeding event.

#### How does mite density affect the burying beetle’s realised thermal niche?

Next, we examined how varying mite density protected burying beetles from competition with blowflies within each temperature treatment (analysing data shown in Fig. 2a-c). We found that beetle breeding success varied with temperature but that the relationship changed with the density of mites (mite x temperature interaction, χ^2^= 138.13, d.f. = 4, *P* < 0.001). To understand the cause of this significant interaction, we split the dataset again into the three different temperature treatments, and compared beetle reproductive success across the three mite treatments. Mites expanded the burying beetle’s thermal niche at lower temperatures. At lower temperatures, mites increased beetle reproductive success, but only at low mite densities. There were more dispersing beetle larvae only when 10 mites were present (*post-hoc* comparison, 0 v. 10 mites: *z* = −3.32, *P* = 0.003) than when 0 mites were present. At intermediate temperatures, beetles had greatest reproductive success when there were 0 (Fig. 2a) and 10 mites (Fig. 2b) present, but suffered a loss in reproductive success when there were 20 mites present (*post-hoc* comparison, 0 v. 20 mites: *z* = 7.76, *P* < 0.001; *post hoc* comparison, 10 v. 20 mites: *z* = 4.87, *P* < 0.001). At higher temperatures, this pattern was reversed, and mites expanded the beetle’s thermal niche. Burying beetles produced more larvae when 20 mites were present than when there were no mites were present at all (*post-hoc* comparison, 0 v. 20 mites: *z* = −7.33, *P* < 0.001).

#### Are mites antagonistic to burying beetles when blowflies are absent?

Here we analysed only the breeding events without blowflies. Temperature affected beetle reproductive success but the effects differed with mite density (mite x temperature interaction, χ^2^= 13.34, d.f. = 4, *P* < 0.010; Fig. 2d-f). To see how, we again split the dataset by temperature. At low temperatures, mites reduced beetle reproductive success, but only at high densities (*post-hoc* comparison 0 v. 20 mites, *z* = 3.95, *P* < 0.001). At intermediate temperatures, mites had a generally negative effect on beetle reproductive success (*post-hoc* comparison 0 v. 10 mites, *z* = 4.23, *P* < 0.001; *post-hoc* comparison 0 v. 20 mites, *z* = 3.93, *P* < 0.001). At higher temperatures, mites again reduced beetle reproductive success, but only at high densities (*post-hoc* comparison 0 v. 20 mites, *z* = 4.19, *P* < 0.001).

#### Do temperature and mite density modulate blowfly success?

Next, we determined the predictors of blowfly reproductive success, analysing only those treatments with blowflies present. We found that the number of blowfly larvae produced varied with temperature, and that this relationship differed with mite density (mite x temperature interaction, χ^2^= 14.33, d.f. = 4, *P* = 0.006; Fig. 3). Splitting the dataset into the three mite treatments, we found that the number of blowfly larvae produced in the absence of mites was substantially greater at higher temperatures than at low (*post-hoc* comparison high v. low temperature: *z* = 3.67, *P* < 0.001) or intermediate temperatures (*post-hoc* comparison high v. mid temperature: *z* = 5.62, *P* < 0.001). Slightly more blowfly larvae were produced at low temperatures than in the mid-temperature treatment (*p*o*st-hoc* comparison low v. mid temperature *z* = 2.56, *P* = 0.029). With 10 mites present, blowfly larvae still thrived at the highest temperatures but performed badly at the lower temperatures (*post-hoc* comparison high v. low temperature: *z* = 4.82, *P* < 0.001; *post-hoc* comparison high v. mid temperature: *z* = 4.33, *P* < 0.001). With 20 mites present, blowflies performed badly at high temperatures (Fig. 3c), and no longer produced more larvae than when breeding at either low (*post-hoc* comparison high v. low temperature: *z* = 1.66, *P* = 0.223) or intermediate temperatures (*post-hoc* comparison high v. intermediate temperature: *z* = 1.27, *P* = 0.415).

**Figure 3.**
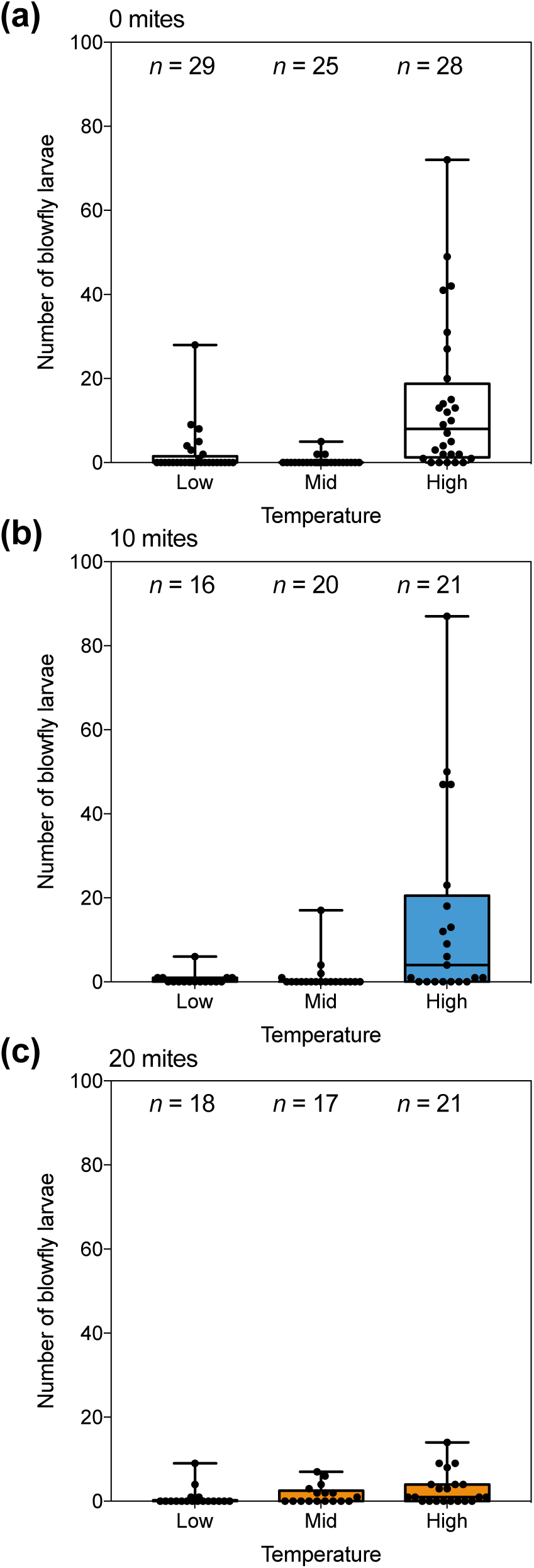
Blowfly reproductive success in relation to temperature in the presence of (a) 0 mites, (b) 10 mites and (c). Sample sizes are as indicated above each bar. The boxplots show the median, inter-quartile range, and the range of data. Each datapoint represents one breeding event.

#### Reproductive success of mites in relation to blowfly presence and temperature

We analysed the effects of blowflies and temperature on mite performance, using two separate analyses, one for the 10 mite treatment and one for the 20 mite treatment. In each case, we found no significant interaction between the blowfly treatment and the temperature treatment on the number of mite deutonymphs produced (10 mites: blowfly x temperature interaction χ^2^= 2.51, d.f. = 2, *P* = 0.286; 20 mites: blowfly x temperature interaction χ^2^= 2.11, d.f. = 2, *P* = 0.349; Fig. S5). Furthermore, when there were 10 mites present we could detect no effect of either temperature (χ^2^= 1.69, d.f. = 2, *P* = 0.429; Fig. S5a,c) or blowfly presence (χ^2^= 0.21, d.f. = 2, *P* = 0.644; Fig. S5a,c) on mite reproductive success. With 20 mites present, there was again no effect of temperature on mite reproductive success (χ^2^= 4.44, d.f. = 2, *P* = 0.108; Fig. S5b,d). The presence of blowflies reduced mite reproductive success (χ^2^= 8.94, d.f. = 2, *P* = 0.003; Fig. S5b,d).

### Laboratory experiment 2

#### Effect of temperature on blowfly larval development

Variation in blowfly survival from egg to the end of larval development could not be explained by the temperature treatment. We placed a similar number of eggs on the mouse at the start of each experiment and found that a similar number of larvae survived to complete development in all three treatments (χ^2^= 0.35, d.f. = 2, *P* = 0.841; Fig. 4a), and that they consumed the carcass to a similar extent (χ^2^= 2.57, d.f. = 2, *P* = 0.277; Fig. 4b).

**Figure 4.**
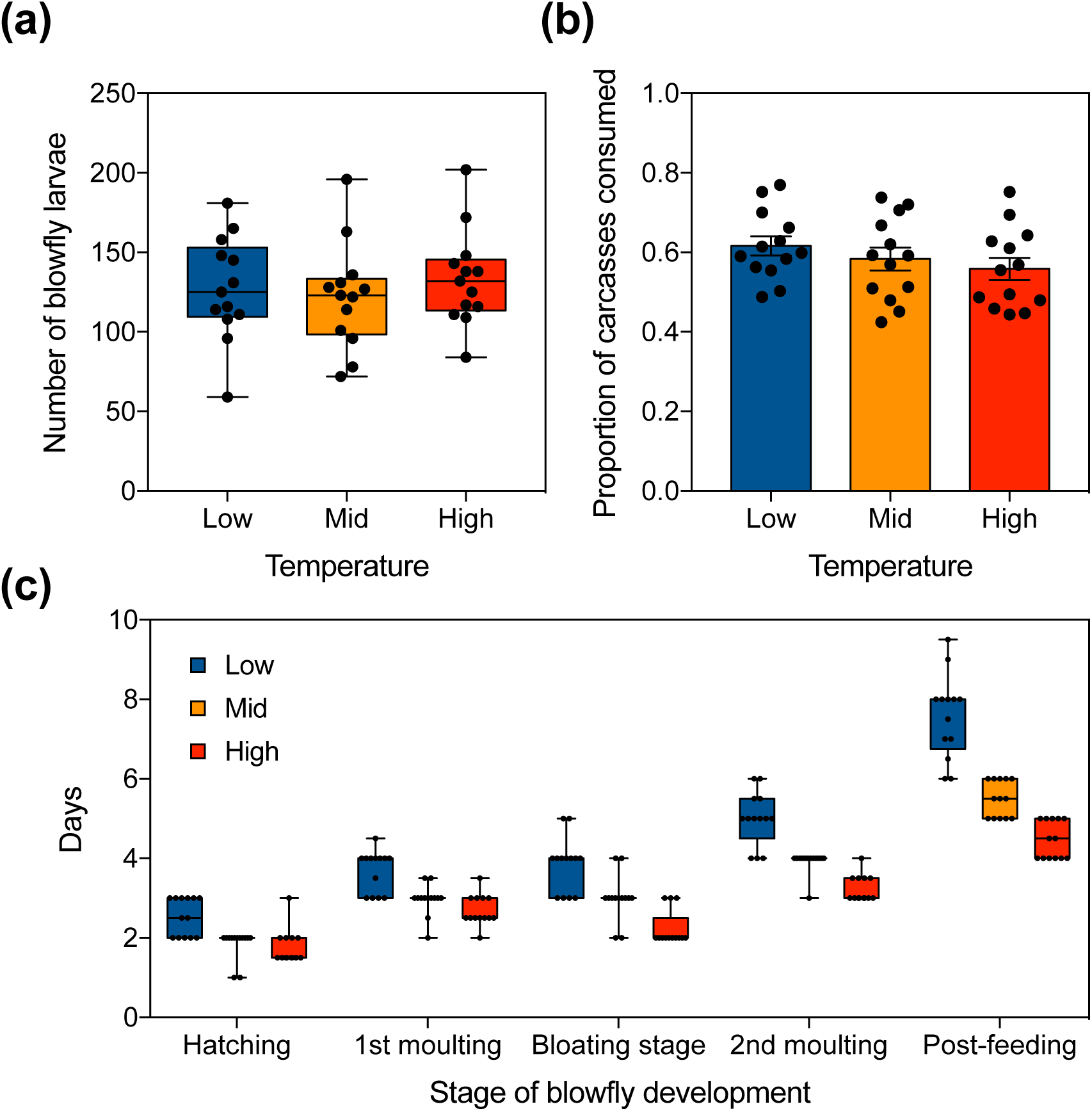
Effect of temperature on different measures of blowfly reproductive performance in relation to temperature. (a) Number of blowfly larvae produced, (b) rate of carcass consumption (mean ± SE), (c) rate of blowfly development. The boxplots show the median, inter-quartile range, and the range of data. Each datapoint represents one breeding event. *n* = 13 mouse carcasses for each temperature treatment.

However, the pace of blowfly development was greatly accelerated at higher temperatures (temperature x developmental stage interaction, χ^2^= 178.46, d.f. = 8, *P* < 0.001; Fig. 4c). Specifically, the first two stages of development, i.e., blowfly eggs and 1^st^ instar larvae were much longer at lower temperatures, compared to development at intermediate (eggs: *t* = 3.75, *P* = 0.001; 1^st^: *t* = 3.93, *P* < 0.001) and higher temperatures (eggs: *t* = −3.76, *P* < 0.001; 1^st^: *t* = −4.89, *P* < 0.001). We could detect no difference in the duration of development between intermediate and high temperatures (eggs: *t* = 0.13, *P* = 0.990; 1^st^: *t* = −0.77, *P* = 0.722). However, carcasses at high temperature reached the bloated stage more rapidly than those at intermediate (*t* = −2.58, *P* = 0.031) and at low temperatures (*t* = −7.33, *P* < 0.001). At high temperatures, blowfly larvae fully consumed carcasses in 4.5 days, whereas larvae at intermediate temperatures took 5.5 days (*t* = −4.39, *P* < 0.001), and those at low temperatures took 7.6 days (*t* = −15.23, *P* < 0.001).

## Discussion

When breeding without blowflies or mites, burying beetles had peak reproductive success at around 15°C, (Fig. 2d: the fundamental thermal niche). Introducing rival blowflies did not change this peak, but caused beetle reproductive success to fall off markedly at lower and high temperatures, both in the field (Fig. 1a) and in the lab (Fig. 2a), reducing the beetle’s realised thermal niche. Our experiments thus suggest that burying beetles can compete effectively, and singlehandedly, with blowflies at 15°C (Fig. 2a,d), perhaps because parents consume blowfly eggs themselves or because beetle larvae are more effective rivals with blowfly larvae when both species develop at this temperature. Whatever the mechanism mediating competition, blowflies showed a corresponding dip in their reproductive success at these intermediate temperatures (Fig. 1d and 3a), which was not seen when blowflies bred alone on a dead mouse (Fig. 4a,b).

Burying beetles were less effective competitors with blowflies at higher and lower temperatures. At higher temperatures, fly development was greatly accelerated (Fig. 4c), making the blowfly larvae more potent rivals for carrion resources. At lower temperatures, burying beetle larvae develop slowly (Meierhofer *et al.* 1999), potentially even more slowly than blowfly larvae (Fig. 4c). This might explain their inferior ability to compete at lower temperatures, but further experiments are needed to test this idea directly.

It was at these higher and lower temperatures that phoretic mites switched from commensalism to a protective mutualism in the field (Fig. 1b,c), causing a substantial reduction in blowfly reproductive success (Fig. 1d-f). The protective mutualism takes the form of pseudo-reciprocity (Connor 2010). The beetle transports the mites to the carcass, and thereby enables the mites to breed, but presumably at some energetic and competitive cost. Nevertheless, by ensuring that mites are present on the carcass, the beetle increases the chance that the mite will return it some fitness benefits during reproduction. Our experiments show that these fitness benefits are derived as a by-product of the mite’s self-serving foraging behaviour rather through a specific adaptation in the mite that has evolved to serve the beetle. We found no evidence that mites are specifically adapted to eat fly eggs because their reproductive success was not enhanced when they could consume fly eggs in addition to carrion (Fig. S5)

The mites expand the burying beetle’s realised thermal niche, counteracting the negative effects of the blowflies. In the field experiments, the burying beetle’s thermal niche extended to include lower temperatures when mites were present at low densities (Fig. 1b) and expanded to include higher temperatures only when mites were present at high densities (Fig. 1c). This might be because blowflies posed the greatest competitive danger to burying beetle larvae at higher temperatures (Fig. 1d), and more mites were required to neutralise this threat. We saw a similar pattern in the protective mutualism when we staged experiments in the lab (Fig. 2b,c). The trapping data show that beetles commonly carry 10-20 mites in the field at higher temperatures, and so are likely to expand the burying beetle’s thermal niche in this way at natural breeding events (Fig S4).

Nevertheless, we found limits on the expression of this protective mutualism. In the lab, high densities of mites significantly decreased beetle reproductive success at intermediate temperatures in the lab, even in the presence of blowflies (Fig. 2c). We did not see equivalent effects under field conditions, perhaps because we made conditions more favourable for blowfly larvae in the lab, by adding them to the carcass well in advance of the beetles and mites. Furthermore, and consistent with previous work (see Wilson 1983) mites were antagonistic to burying beetles at all temperatures unless blowflies are present (Fig. 2e,f).

Previous studies have emphasised the significance of the abiotic environment in tipping interactions from mutualism to parasitism (Chamberlain *et al.* 2014; Hoeksema & Bruna 2015; Gorter *et al.* 2016). For example, protective mutualisms sometimes break down at higher temperatures because the protecting partner is the more vulnerable to heat stress when temperatures rise (e.g. Barton & Ives 2014; Fitzpatrick *et al.* 2014; Doremus & Oliver 2017). However, we found no evidence that mites were more vulnerable to higher temperatures, at least under field conditions. Instead, the main driver of change in the protective mutualism came from the response of enemy blowflies to variation in temperature. We suggest that similar effects might be found in other protective mutualisms, providing that both partners can tolerate some thermal stress, and where enemy species are more likely to thrive at high temperatures. Predicting how a species might respond to climate change thus involves understanding its interactions within the natural ecological community as well as some knowledge of the intrinsic variation among those interacting species in their thermal tolerance (Early & Keith 2019).

In conclusion, our experiments show that not all mutualisms are vulnerable to collapse under rising temperatures (see also Frederickson 2017). Where both partners have a similar degree of thermal tolerance, mutualisms can instead play a key role in ensuring that species persist in new and adverse thermal environments (Afkhami *et al.* 2014; Johnson 2015; Peay 2016), by expanding the realised thermal niche.

## Acknowledgements

This project was supported by the Rosemary Grant Award from the Society for the Study of Evolution. S.-J.S. was supported by the Taiwan Cambridge Scholarship from the Cambridge Commonwealth, European & International Trust. R.M.K. was supported by a European Research Council Consolidator grant 301785 BALDWINIAN_BEETLES and a Wolfson Merit Award from the Royal Society.

## Supplementary Information

### Maintenance of the colonies of beetles, mites, blowflies

*Burying beetles* We bred burying beetles by introducing pairs of unrelated males and females to a mouse carcass (7-15 g) in a plastic container (17 × 12 × 6 cm filled with 2 cm of moist soil). All larvae were collected at dispersal, and transferred to eclosion boxes (10 × 10 × 2 cm, 25 compartments) filled with damp soil. Once they had developed into adults, beetles were kept individually in plastic containers (12 × 8 × 2 cm) filled with moist soil, being fed twice a week with small pieces of minced beef.

*Mites* We maintained mite colonies in plastic containers (17 × 12 × 6 cm filled with 2 cm of moist soil). Each container was provided with an adult beetle and fed with pieces of minced beef twice a week. We bred mites once a month by introducing 15 mite deutonymphs to a pair of beetles and a mouse carcass in plastic containers (17 × 12 × 6 cm filled with 2 cm of moist soil; *n* = 10). When the burying beetle larvae had completed their development, we collected mite deutonymphs that were dispersing on adult beetles. Newly-emerged mites were reintroduced to the containers holding the mite colony.

*Blowflies* Colonies of blowflies (*n* = 5) were reared in screened cages (32.5 × 32.5 × 32.5 cm. They were continuously supplied with a mixture of powdered milk and dry granulated sugar, and ad lib. water. We fed newly emerged blowflies with pig livers to induce maturation of the ovaries. After a week, these blowflies were then given mouse carcasses to breed upon. All beetle, mite, and blowfly colonies were kept at 21 ± 2°C with a photoperiod of 16:8 light:dark.

## Statistical Methods

### Burying beetles and phoretic mites in Madingley Wood

We analysed how the number of beetles collected from each trap, and the number of mites borne by each individual beetle, varied with temperature, using a separate GLMM for each measure and a negative binomial distribution to account for data over-dispersion. Each trapping event was treated as an independent datapoint. In each model, we included temperature and its quadratic term as covariates, to account for any potentially non-linear effects of temperature on beetle or mite abundance. Temperature was measured as averaged daily mean temperature during the two week period in which the trap was set, starting from trap setting to beetle collection. Trap ID and year were included as random factors. In the model with mite number per beetle as the independent variable, we also included beetle sex as an explanatory factor.

### Field experiments

We sought correlates of beetle brood size, the number of blowfly larvae, and the number of mite offspring number at the end of each trial, using three separate GLMMs each with negative binomial distributions. For the models with beetle brood size and the number of blowfly larvae as independent variables, we included carcass mass, mite treatment, temperature, and the interaction between mite treatment and temperature as covariates. Temperature was calculated as the average daily mean temperature, from carcass introduction to larvae dispersal (or carcass consumption by blowfly larvae). We also included a squared measure of temperature in the model to allow for any possible potential non-linear relationships. The model analysing mite reproductive success included data from the treatments with 10 and 20 mites and included carcass mass and temperature as covariates. In all three models, experimental site and year were included as random factors.

### Laboratory experiments

*Analyses of beetle reproductive success* We tested for the interacting effects of blowfly, mite, and temperature treatments on the reproductive success of beetles, using GLMMs, including all three treatments as categorical covariates, and using a Poisson distribution. We split the dataset by treatments to determine how any significant interactions arose.

*Analyses of blowfly reproductive success* We tested for the interacting effects of mites and temperature using GLMMs, using a negative binomial distribution to account for data overdispersion, and again including these two effects as categorical covariates. We split the dataset by treatments to determine how any significant interactions arose.

*Analyses of mite reproductive success* We tested how mite reproductive success covaried with the temperature treatment using a GLMM with a negative binomial distribution to account for data overdispersion. Using a similar model, we further compared their reproductive success in the presence and absence of blowflies. In all three types of model, we included block as a random factor.

#### Effect of temperature on blowfly larval development

We analysed the number of blowfly larvae in a negative binomial regression model with the function *glm.nb* in the MASS package to account for overdispersion. We analysed carcass consumption rate in a beta regression model in the *betareg* package. In both analyses, we included temperature treatment (low, intermediate, and high), blowfly egg mass, and carcass mass as covariates. To analyse the effect of temperature on the developmental rate of blowfly larvae, we used a GLMM with Gaussian error structure and included the interaction between temperature treatment and developmental stage, blowfly egg mass, and carcass mass as covariates. In this analysis, we also included the ID of each carcass as a random factor, since carcasses were sampled repeatedly across different developmental stages.

## Supplementary Figures

**Figure S1.**
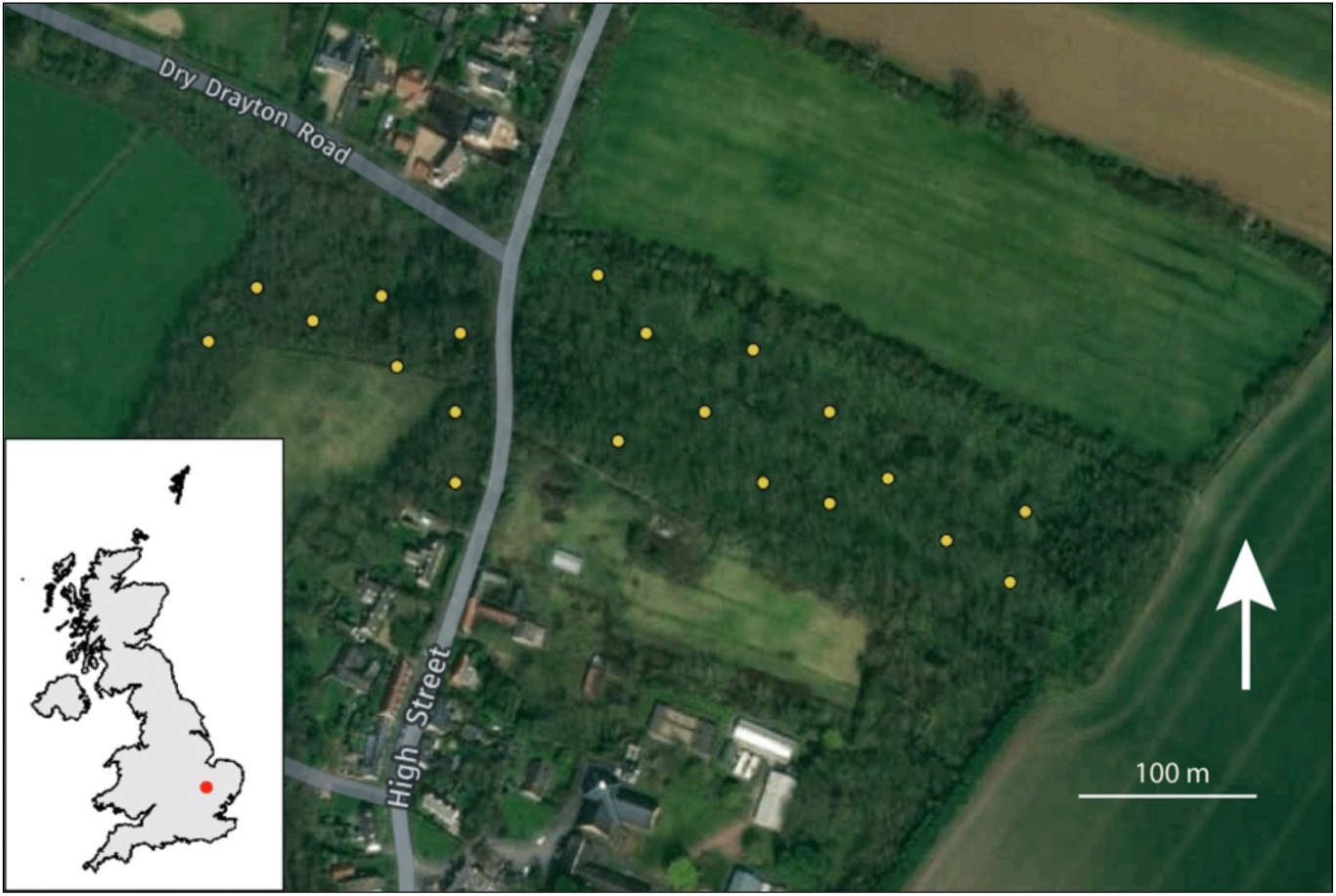
Spatial distribution of breeding sites (yellow dots) established in the field experiment at the study in Madingley Wood, Cambridge, UK (Latitude: 52.22730°; Longitude: 0.04442°). Image taken from GoogleMaps.

**Figure S2.**
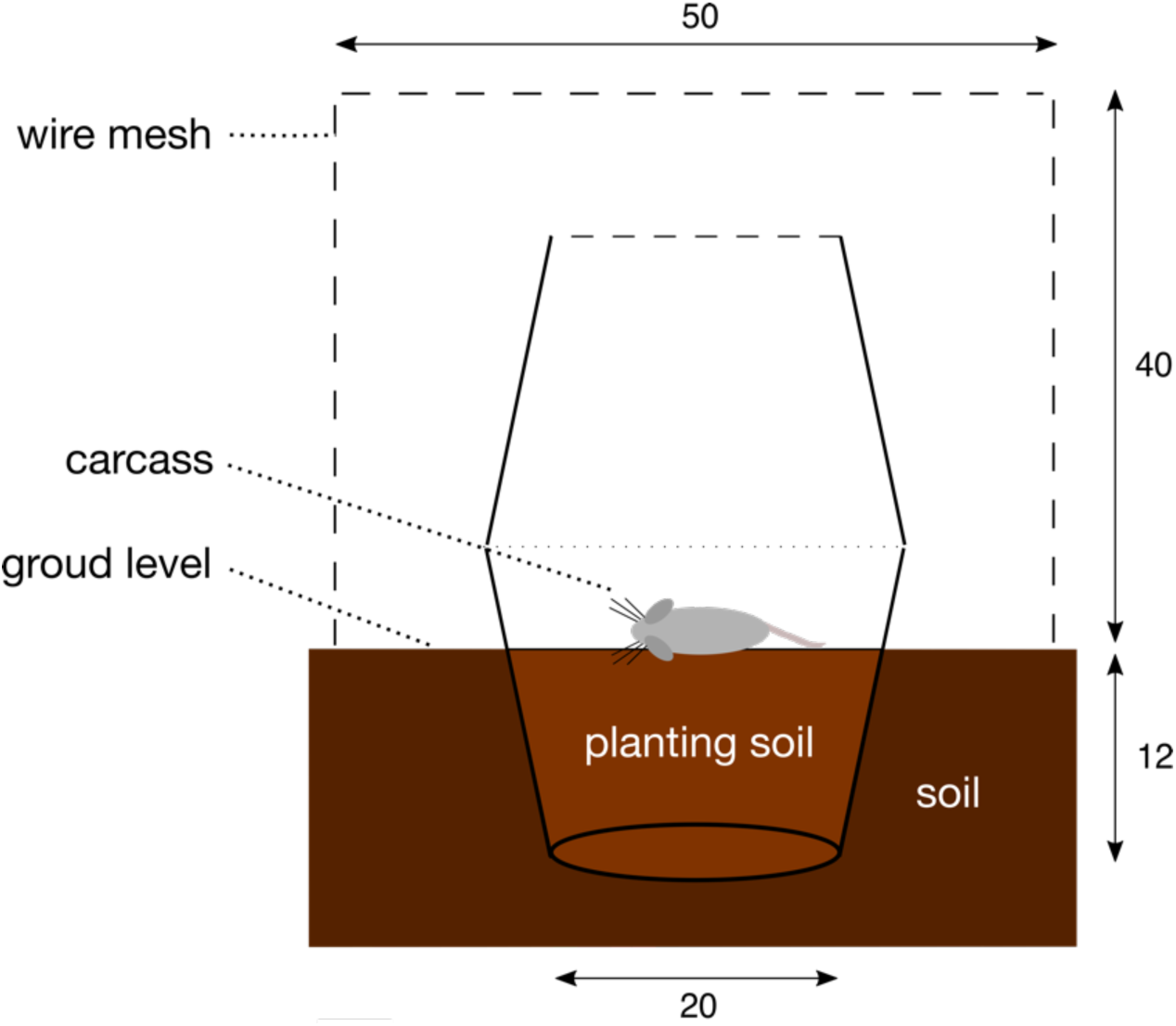
Schematic side-view representation of the experimental setup used for each breeding event in the field (dimensions are in cm). One flowerpot was partially buried in the ground, filled with compost (planting soil) and covered above with a second inverted flowerpot, perforated on the top to let in blowflies. The whole apparatus was surrounded by wire mesh, pegged in the ground, to prevent disruption by scavengers.

**Figure S3.**
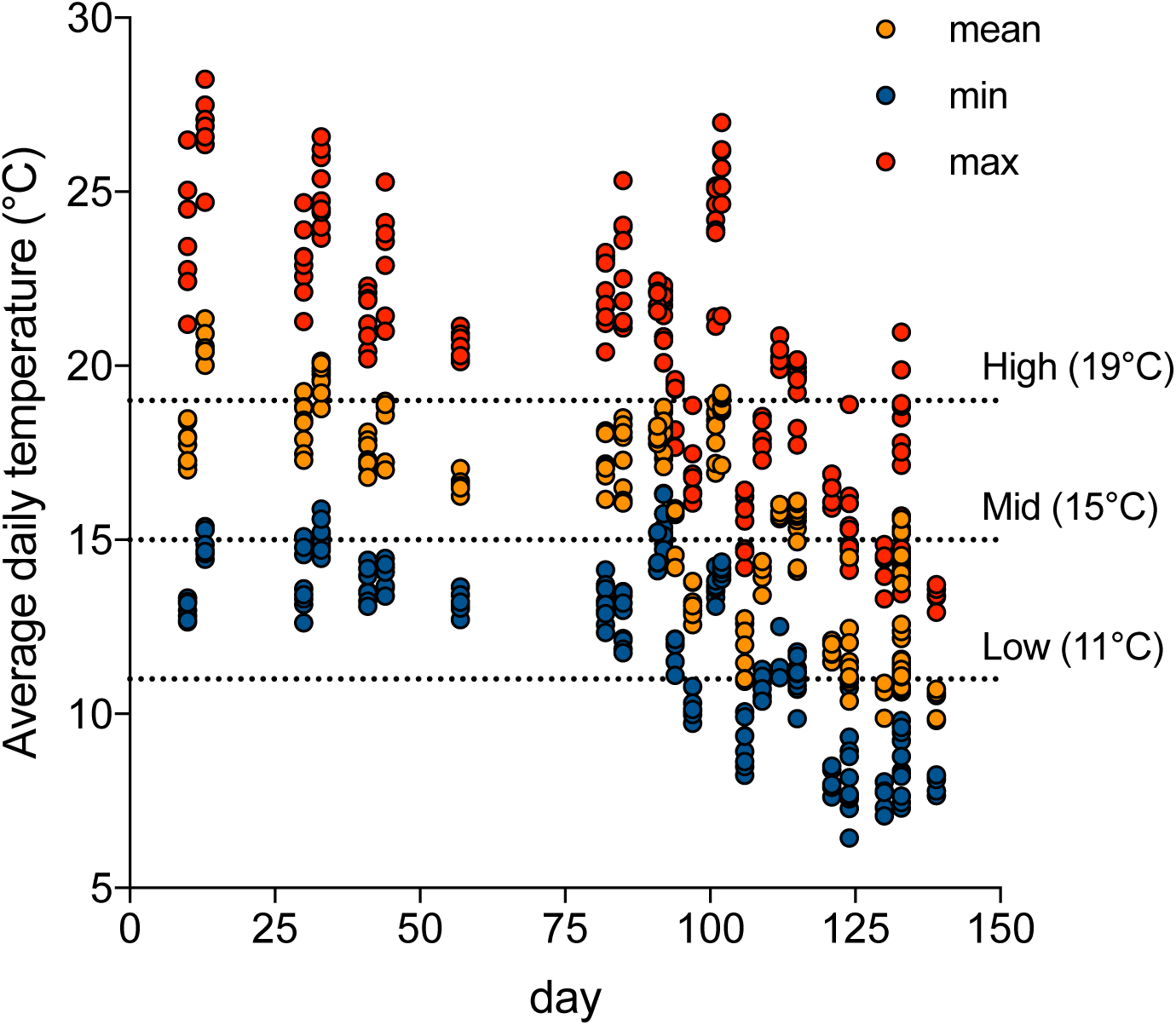
Daily variation in the average daily mean, maximum, and minimum ambient air temperature in Madingley Woods during the field experiments conducted in 2016 and 2017. Day 0 is June 1. Dashed lines correspond to the high, mid, and low temperatures used in the laboratory experiments.

**Figure S4.**
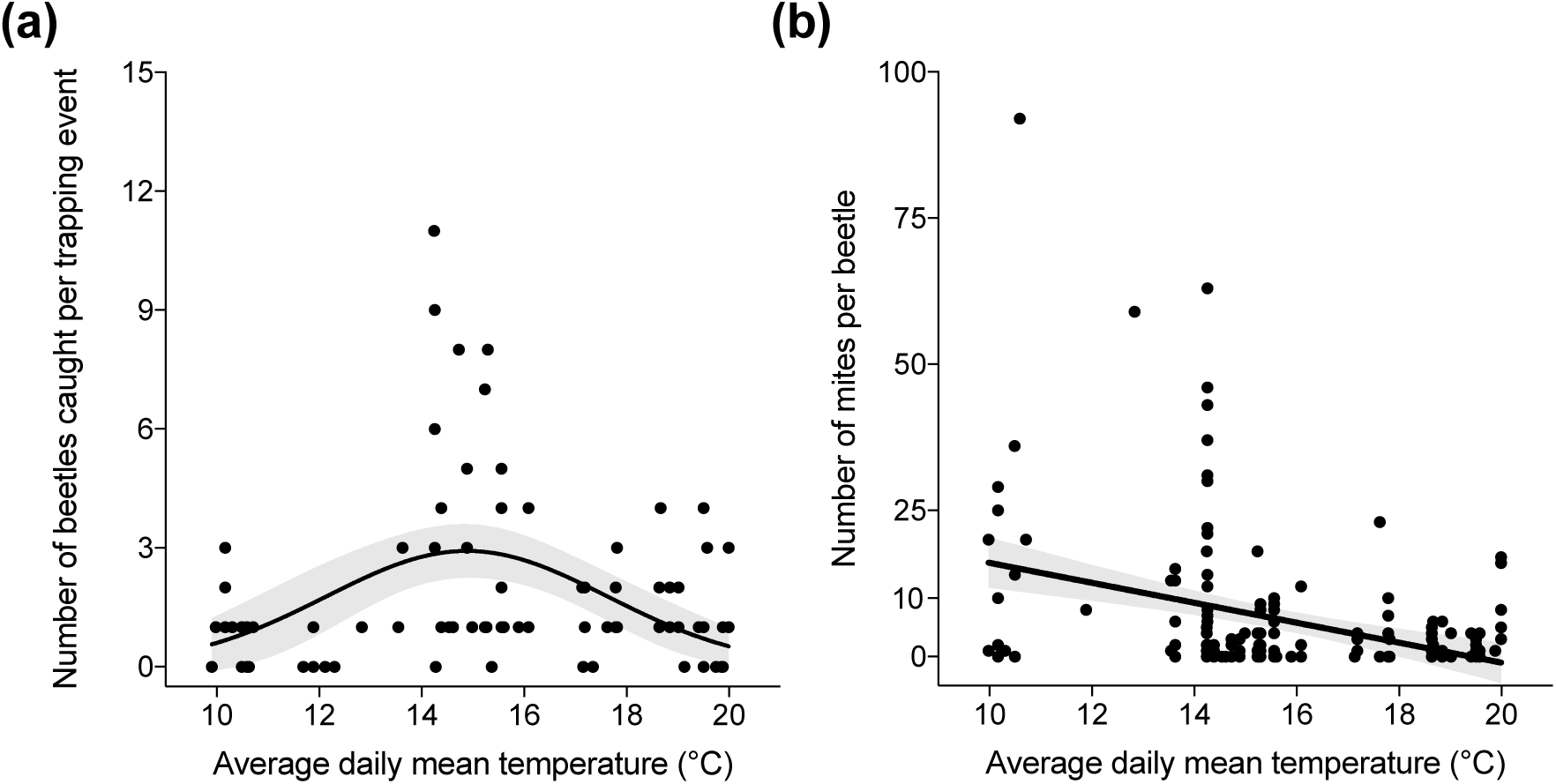
Relationship between ambient air temperature in Madingley Woods and number of *N. vespilloides* caught per trapping event (*n* = 98 trapping events) and the number of mites carried by each beetle (*n* = 172 beetles) at each trapping event. The lines denote predicted relationship in GLMM, and shaded areas represent 95% confidence intervals.

**Figure S5.**
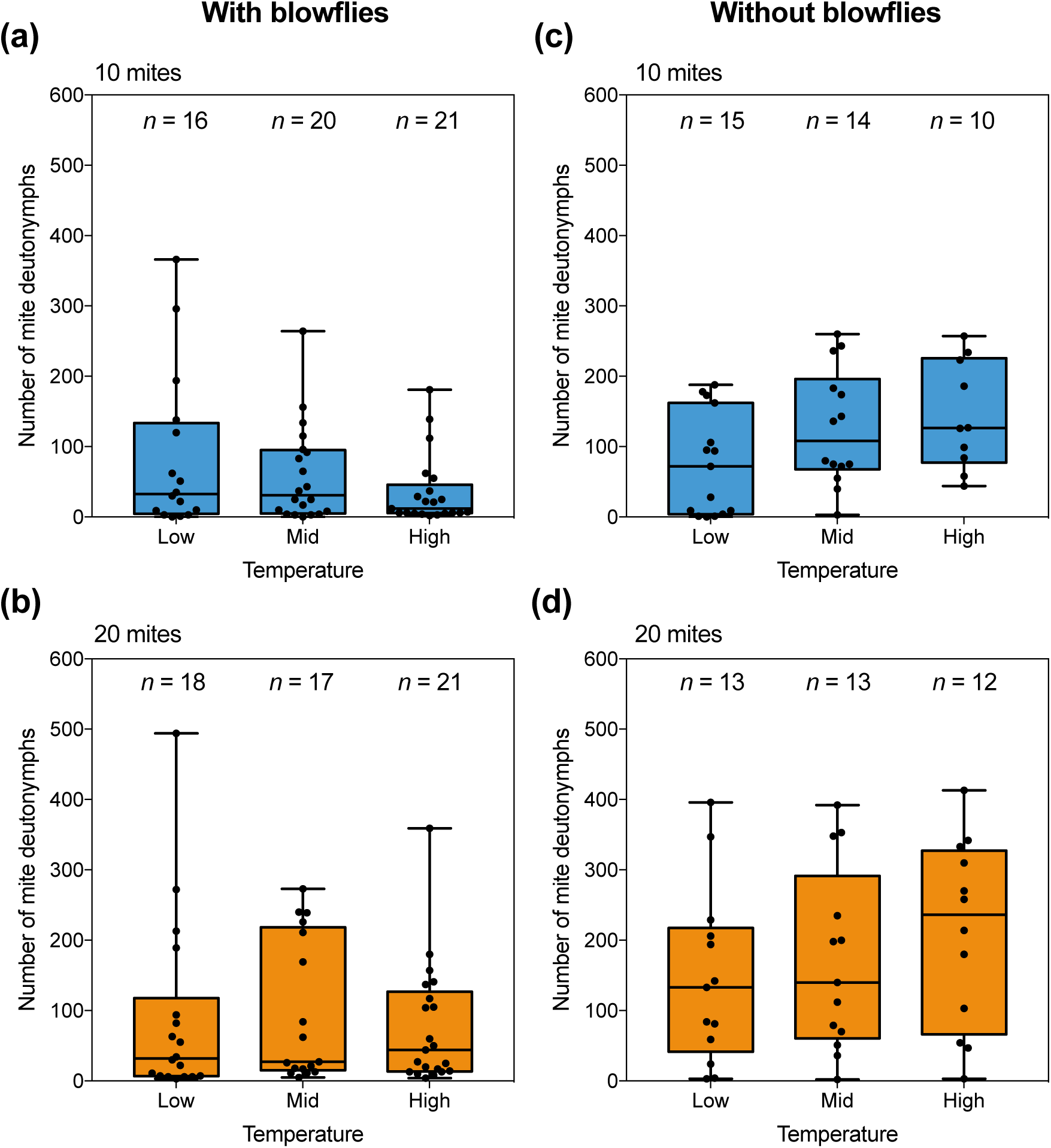
Reproductive success of mites in relation to temperature, with and without blowflies and across the temperature treatments. Data for each mite treatment (10 and 20 mites) are shown separately. Sample sizes are as indicated above each boxplot. The boxplots show the median, inter-quartile range, and the range of data. Each datapoint represents one breeding event.

